# Self-activated photoblinking of nitrogen vacancy centers in nanodiamonds (sandSTORM): A method for rapid single molecule localization microscopy with unlimited observation time

**DOI:** 10.1101/2020.05.20.106716

**Authors:** Kaarjel K. Narayanasamy, Joshua C. Price, Raquel Mesquita-Riberio, Melissa L. Mather, Izzy Jayasinghe

## Abstract

Stochastic optical reconstruction microscopy (STORM) is one of the most commonly used super-resolution microscopy techniques. Popular implementations of STORM utilize aromatic fluorophores and consist of a number of intrinsic limitations such the finite photostability of the dyes, the reliance upon non-physiological redox buffers and speed which is ultimately limited by the ‘off’-rates of the photoblinking. Self-activated nanodiamond-based STORM (sandSTORM) has been developed as an accelerated STORM protocol which harvests the rapid, high quantum-yield and sustained photoblinking of nanodiamonds (ND). Photoluminescence emanating from the stochastic charge-state interconversion of Nitrogen Vacancy (NV) centers between NV0and NV^-^ is localized using conventional STORM-optimized hardware and image processing protocols over an unlimited duration of imaging. This produces super-resolution images of matching resolution at ∼ 3-times the speed and ∼ 100 times less light exposure to the sample compared to traditional STORM. The enabling NDs have been used to map arrays of ryanodine receptor in skeletal muscle tissues via immunolabelling and directly visualize the internal spaces of living neurons via endocytosis of NDs. This paper details the physical basis of sandSTORM, factors which optimize its performance, and key characteristics which make it a powerful STORM protocol suitable for imaging nanoscale sub-cellular structures.

## Introduction

Super-resolution microscopy has transformed cell and molecular biology by breaking the diffraction limit of light enabling biological structures critical to cell function and life to be resolved at a molecular scale. Commonly-visualized structures include cellular organelles [1, 2], cytoskeleton [3, 4], nucleic acid organization [5, 6] and protein clustering [7, 8]; see review [9]. Among many super-resolution techniques which continue to emerge and evolve, single molecule localization microscopy (SMLM) is arguably the most popular modality used by life scientists, with the variant known as STORM widely used, largely owing to its compatibility with well-established fluorescent marker probes and immuno-labelling protocols [10-12]. The basis of STORM is the separation of consecutively emitting fluorescent sources within diffraction-limited spots from sequential images. Post processing is performed to localize fluorescent reporters within a single reconstructed image using centroid-localization algorithms and Gaussian fitting methods achieving precision down to ≤ 20 nm [13]. Thousands of images are typically acquired to improve localization precision based on activation of stochastically different fluorescent emitters within diffraction limited spots [14].

A fundamentally important element of attaining high quality super resolution images via STORM is the use of fluorescent probes that exhibit at least two states that last for sufficient time to enable these to be distinguished as ‘on’ and ‘off’ [14]. Fluorescent probes typically used in STORM include organic fluorophores, fluorescent proteins and solid-state nanoparticles. The photophysical behavior of such probes can be controlled via multiple mechanisms including light induced interconversion between different spectroscopic states [11, 15], chemically mediated blinking [16, 17], reversible photoisomerization of chemical groups [18] and transient binding and unbinding of probes [19, 20]. Fluorescent probe selection and design requires consideration of probe brightness and the ratio between the brightness of the probe in the ‘on’ and ‘off’ states, as these parameters have a significant impact on the localization precision [14]. The photostability, number of cycles over which the probe can change state and the switching duty cycle, are also important factors affecting image quality, acquisition time and the available measurement window.

Efforts to improve fluorescent probes for localization microscopy are ongoing. Recent refinements tackle issues such as the availability of switchable probes, integration of protocols with established labelling methods, the rapid photobleaching of organic probes and the intrinsically poor temporal resolution of such imaging experiments which owes to the need to acquire thousands of images. Many of these shortcomings have been addressed using direct STORM (dSTORM) which is a method that enables the use of conventional fluorescent probes such as labelled antibodies or chemical dyes (e.g. cyanine dyes) without the need for additional activator fluorophores, required in previous methodologies [17].

Research is also underway to address the limited quantitative capacity of STORM arising as a result of the lack of control over fluorophore stochastic photo-switching and photo-bleaching [16, 21, 22] in addition to the inherent slowness of the imaging modality. Further advances in STORM have been made through the implementation of alternative probes based on solid-state nanoparticles such as pure metal nanoparticles [23], lanthanide-doped nanoparticles [24], carbon dots [25] and quantum dots [26]. Advantageously these probes have been demonstrated to address the rapid photobleaching which occurs in organic fluorophores opening the door to time course and repeated studies.

Currently, STORM is more commonly implemented using fixed cells however there is tremendous interest in establishing STORM techniques that enable live cell time course imaging. This demands fast cycling fluorescent probes that are resistant to photobleaching, produce high contrast between the ‘on’ and ‘off’ states, and in the case of intracellular imaging can be transported across the cell membrane. The realization of this will reduce the number of images required for reconstruction and increase the achievable frame rate, minimizing motional artefacts and increasing sample throughput. Typical STORM imaging times for fixed samples are of the order of 20-30 minutes and to meet the demands of live cell imaging and/or high-throughput imaging, these times need to be reduced by at least an order of magnitude. Allied to work centered on fluorescent probe optimization, are studies seeking to increase imaging speeds based on algorithmic enhancement of the image reconstruction and through the establishment of algorithms to accommodate either high [27, 28] or sparse marker localization densities [29]. Conspicuously lacking however, have been experimental approaches which offer genuine acceleration of STORM through a reduction of the time between localizations (i.e. ‘off’ times) without compromising the resolution or the durability of the samples.

Herein a method for rapid SMLM with unlimited observation time based on self-activated photoblinking of Nitrogen Vacancy (NV) centers in nanosized diamond (nanodiamonds (ND)) is reported. The NV center is a naturally occurring paramagnetic impurity comprising a substitutional Nitrogen atom adjacent to a vacant site in the diamond lattice. This defect has been reported to exist in three different charge states namely the negative charge state (NV^-^), neutral charge state (NV°) and positive charge state (NV^+^), defined by the number of unpaired electrons nearby the defect. The NV^-^ center in particular has attracted attention as a potential fluorescent probe for use in biological imaging due to its high quantum yield, robust luminescence, which does not bleach, and the inherent low cytotoxicity of diamond [30, 31]. The method presented here leverages the intrinsically-driven interconversion between the NVoand NV^-^ charges states mediated via photoionization and the surrounding chemical environment [31]. Photoluminescence of NV centers has been previously been used for deterministic super-resolution techniques such as Stimulated Emission Depletion (STED) and Ground State Depletion (GSD) microscopies [15, 32, 33]. To use them as probes for localization microscopy, it has always been necessary to manipulate the spin transitions of NV centres with the use of extrinsic microwave which has limited their utility in imaging biological nanostructures [30, 34-36]. The present work is the first time, to the authors’ knowledge, SMLM based on the *stochastic* blinking of NVs has been demonstrated on biological samples. In particular, the intracellular ryanodine receptor (RyR) ion channel is imaged within skeletal muscle tissue using antibody conjugated NDs containing NVs. Additionally, endocytosed nanodiamonds are localized with super-resolution in networks of neuronal cells. Further, this work demonstrates how NVs can be photo-switched at very short dark times by carefully controlling the emission wavelength used for localization and the electrolyte solution samples are immersed in. Significantly, sandSTORM is shown to enable up to 10 times higher frames rates (100 fps compared with 10-20 fps) than dSTORM with the added advantages of simpler probe preparation, compatibility with basic cell culture buffers and use of probes that are resistant to photobleaching.

## Results and Discussion

### sandSTORM

#### Super-resolution localization of ion channel arrays

Time-sequential wide field images of transverse cryosections of extensor digitorum longus (EDL) muscles from rats were acquired based on the photoluminescence from NVs within commercially sourced NDs co-localized with RyR calcium release channels. Experimentally NVs were excited using a 561 nm laser line and the resulting photoluminescence within a 605 nm ± 35 nm pass band used to form individual image frames. Post processing of ∼ 30,000 images was subsequently performed to reconstruct the RyR array networks with super-resolution based on the use of self-activated photoblinking of NVs within proximal NDs, the results of which are shown in Figure 1. The architecture typical of RyRs in transversely sectioned skeletal muscle fibers is faithfully depicted through a comparison with a corresponding diffraction limited wide-field fluorescent image (Figure 1 inset). The enhanced localization sandSTORM produces is apparent and the nanoscale morphology that it reports of the RyR arrays reproduces the observations of previous investigations using standard dSTORM [37].

**Fig. 1.**
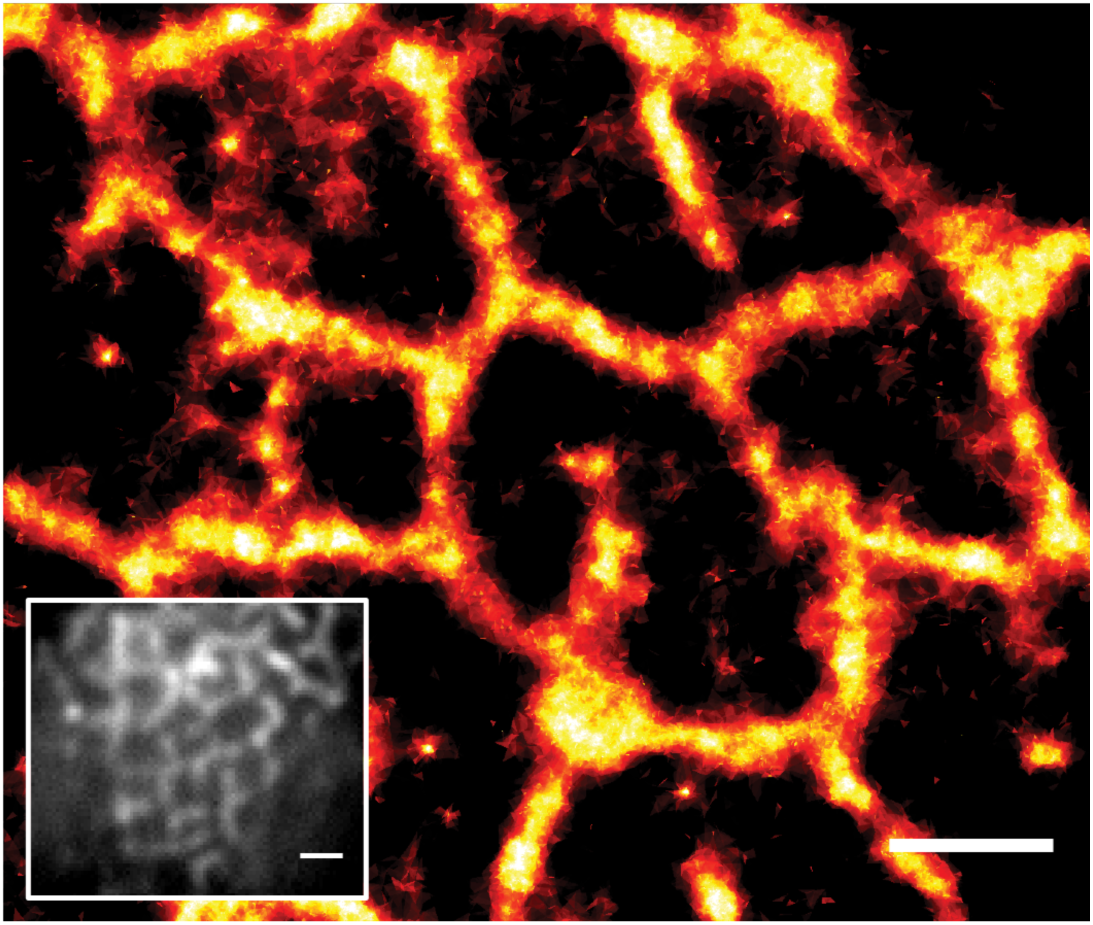
sandSTORM image of RyRs labelled with ND-conjugated antibodies in a rat skeletal muscle transverse tissue section (scale bar = 1 µm). Inset displays diffraction limited image of an equivalent region (scale bar = 2 µm)

The ability to resolve RyR arrays based on photoblinking from NVs using different spectral ranges of emitted light was also investigated based on the hypothesis that photoblinking is linked to NV charge state conversion between the NV° and NV^-^ states. These charge states have intrinsically different electronic structures and correspondingly different emission spectra [38], which are characterized by narrow zero phonon lines at 575 nm and 637 nm for the NV° and NV^-^ charge states respectively and broad phonon lines extending as far as 750 nm for NV° and 800 nm for NV^-^. Experimentally the contribution from each charge state on the reconstructed sandSTORM image was investigated through the use of five different emission filters (520 nm ± 20 nm bandpass (BP), 575 nm ± 2.5 nm BP, 590 nm longpass (LP), 605 nm ± 35 nm BP, 692 nm ± 20 nm BP) covering spectral ranges that selectively include or exclude contributions from the individual charge states. Figure 2 displays exemplar unprocessed images indicative of the photoluminescence detected using each emission filter (top row) and the resulting reconstructed sandSTORM images (bottom row).

**Fig. 2.**
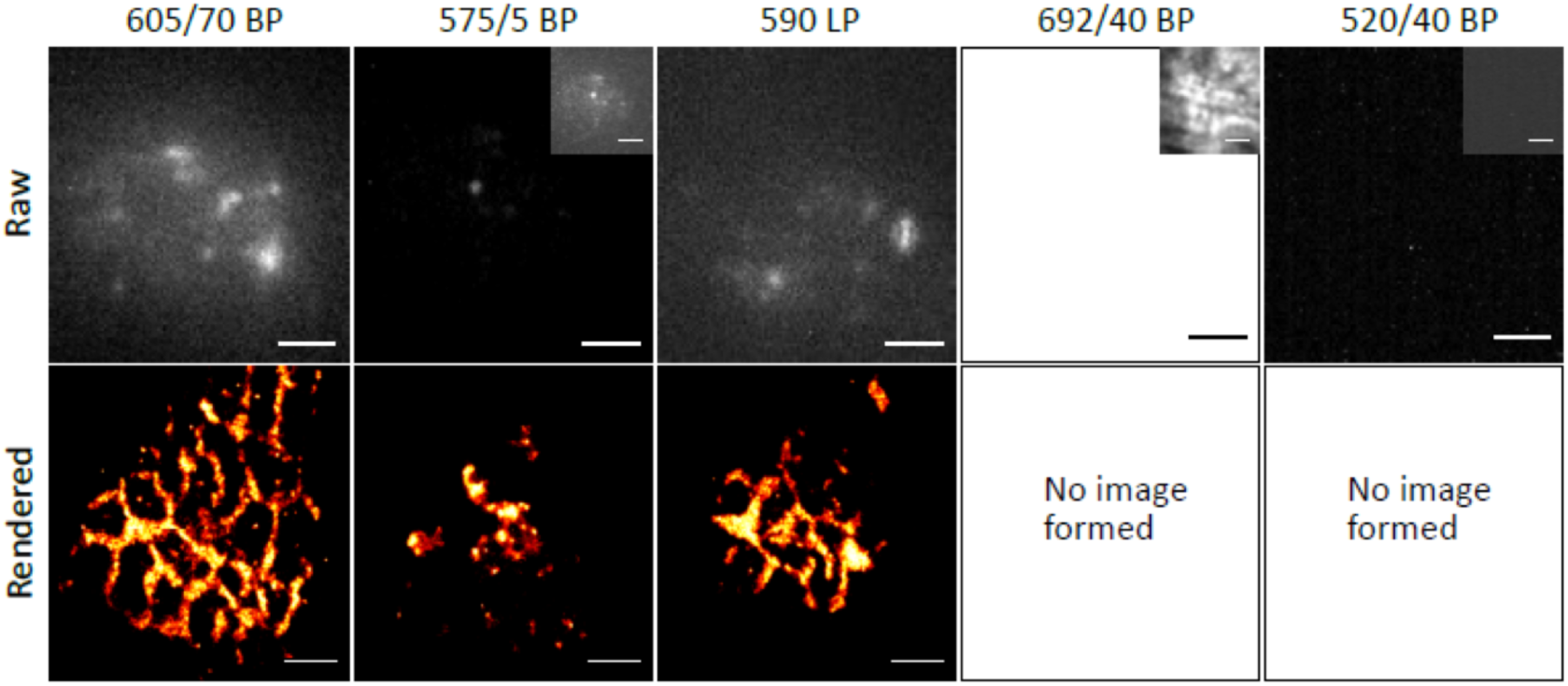
The spectral band-specific reconstruction of sandSTORM images. Exemplar individual image frames (top row, inserts show images with enhanced contrast) and corresponding sandSTORM reconstructions (bottom row) from experiments carried out using five different emission filters (left to right: 605 nm ± 35 nm BP, 575 nm ± 2.5 nm BP, 590 nm LP, 692 nm ± 20 nm BP, 520 nm ± 20 nm BP). Scale bar = 2 µm.

Photoblinking was observed to be most prominent in the spectral band in which NVoand NV^-^ emission overlaps (570 nm to 640 nm). There was either insufficient blinking and/or inadequate contrast between the ‘on’ and ‘off’ states outside of these bands for sandSTORM images to be formed. Spectral windows extending well beyond these bands (e.g. 590 LP filter, Figure 2) also featured diminishing contrast in the raw images which translated either to poor detection of photoblinks or worse localization error. With the sandSTORM experimental protocol used here, effective localization requires collection of light from a mixed population of charge states in a spectral region in which the NV^-^ emission does not dominate over NV°. Using the 520/40 BP emission filter, corresponding to a region outside of the NV° and NV^-^ emission band no photoblinking was observed.

This spectral dependence of sandSTORM image reconstruction suggests the mechanisms for photoblinking are transient conversions between the NV° and NV^-^ charge states. Owing to the light intensity used in this work, 1.04×10^6^ W cm^-2^, photoinduced electron transfer is thought to be the main driver of charge state conversion and correspondingly photoblinking. Indeed, previous reports [39-41] demonstrate a photoinduced switch from the NV° to the NV^-^ state that was linked to ionization of Nitrogen donors proximal to NVs which involved electron transfer from nearby substitutional N was to the NV centre. Further support for this hypothesis is work on the photophysics of the NV that has shown the defects’ charge state is dynamically modulated between NVoand NV^-^ during illumination with green light by Aslam, Waldherr, Neumann, Jelezko and Wrachtrup [40]. Specifically, their work looked at the dynamics of charge carrier diffusion and trapping that demonstrated efficient production of holes from green laser induced conversion between NVoand NV^-^. Their findings demonstrate the charge state of NVs is stable in the presence of free electrons and by observing the distribution of trapped charge after local photoionization NVs were found to be effective hole traps, resistant to conduction electrons. They considered the NV^-^ hole capture rate and the NV° electron capture rate in terms of the charge state conversion rate [42]. An additional mechanism causing photoblinking includes NV^-^ ionization into NV° which is known to depend on the laser power used and the excitation wavelength. Direct ionization of NV^-^ requires photon energies higher than 2.6 eV which is not possible using the 561 nm illumination in the work here, corresponding to, 2.21 eV. However, ionization may occur via a 2 photon absorption process in which one photon induces a transition to the excited state of the defect while the second photon excites the electron to the conduction band of the diamond [43]. Blinking on the microsecond time scale has been ascribed to a photoconversion process of the NV itself between the NV^-^ and NV° state whilst blinking occurring at the millisecond time scale has been linked with a two photon charge state conversion process. Although it is difficult to elucidate the exact mechanism(s) driving photoblinking in the work herein, photoionization leading to conversion from the NV to the NV-charge states are thought to dominate.

#### Super-resolution localization of NDs within neuronal cells

To further demonstrate the use of sandSTORM in biological systems endocytosed NDs within networks of cortical neuronal cells were imaged, Figure 3. Experimentally NDs were introduced into live cultures of cortical neuronal cells following 14 days of culture and incubated for 24 hours to allow cell mediated endocytosis of NDs to occur, as depicted in Figure 3a. Cells were subsequently fixed for performing counter-stain immunolabelling to identify cell nuclei and axons and the relative localization of NDs to be visualized (Figure 3b). Fixation was optional as sandSTORM with internalized NDs was compatible with the living culture maintained in PBS. Diffraction limited (Figure 3c) and sandSTORM (Figure 3d) images of NDs were obtained which depict the structural organization of the thread-like structures of axons within the neuronal network, indicating co-localization of NDs with axons. The enhanced localization achieved with sandSTORM as compared to the diffraction limited fluorescent image is apparent and additionally confirms the photoblinking capacity of NVs is retained once internalized in cells. These results further demonstrate that, in addition to targeting the NDs to protein targets via antibodies, they can be directly used to monitor both endocytotic pathways (with potential utility in visualizing targeted drug delivery) and visualize nanoscale intracellular spaces within living cells.

**Fig. 3.**
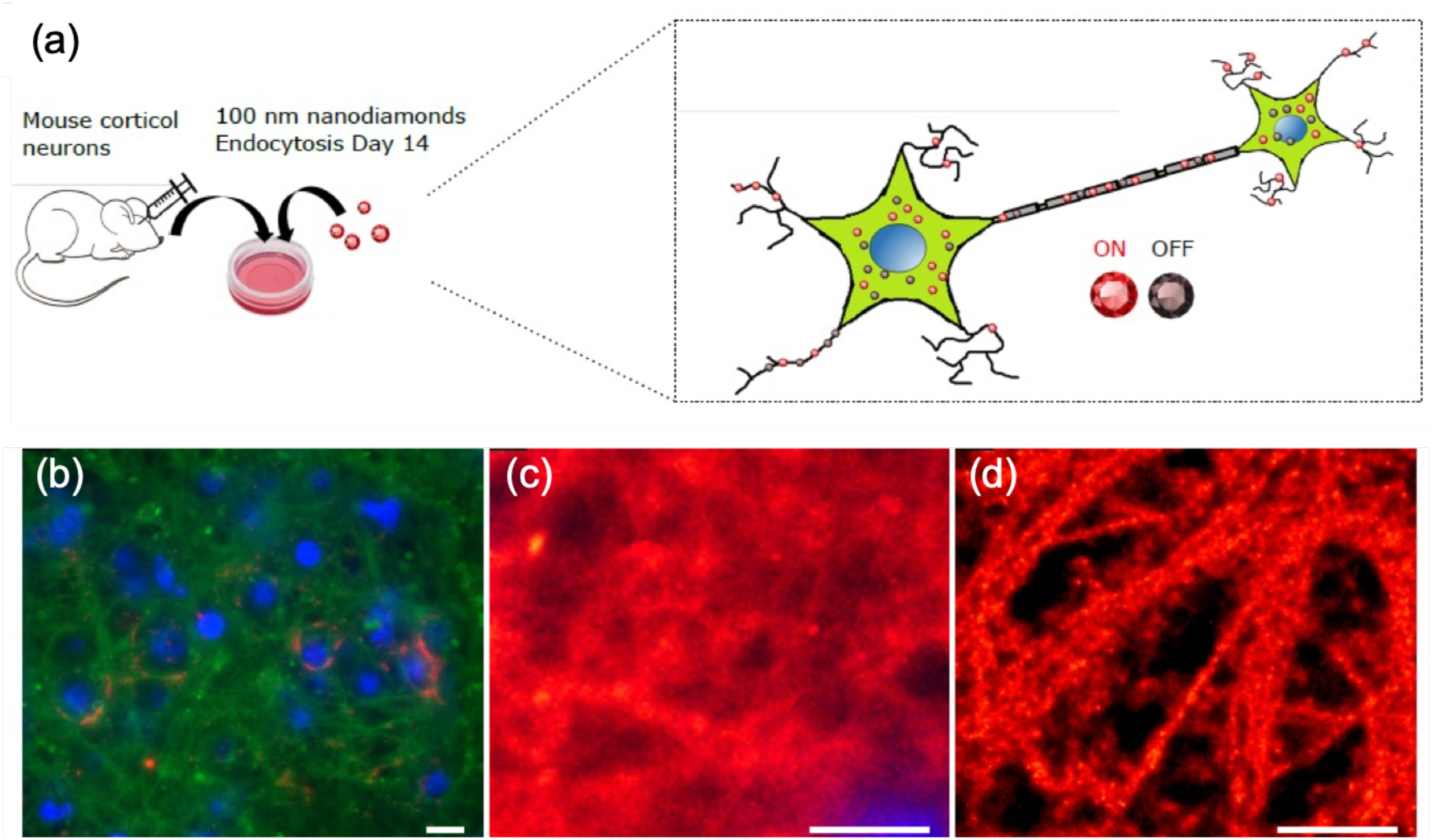
Imaging internal spaces of cultured neurons with sandSTORM. (a) Schematic of experimental protocol for incorporation of NDs within live neuronal cells. (b) corresponding immunofluorescence, (c) diffraction limited (scale bar 10 µm) and (d) sandSTORM (scale bar 2 µm) (d) images of fixed cells (scale bar 2 µm).

#### NV blinking in different electrolyte solutions

Factors affecting NV photoblinking were investigated with an emphasis on the chemical environment local to NDs. Herein, spatially localized blinking events were tracked from NDs which have labelled RyR in a muscle tissue section (similar to region shown in Figure 1), immersed sequentially in five different electrolyte aqueous solutions (Hydrochloric acid (HCl), acetic acid (CH_3_COOH), sodium peroxide (NaOH), sodium chloride (NaCl), phosphate buffered saline (PBS)). Figure 4a displays a subset of time course data for each electrolyte solution studied representative of the number of blinking events recorded per image frame (40,000 frames, 10 ms exposure time, 5 repeats), extracted using a threshold method (Supplementary Figure 1). The results indicate a dependence of photoblinking on the electrolyte solution in terms of both the number of blinking events per frame and the frequency of events over time (see also Supplementary Movie). To investigate this further, the blinking on and off times for each solution were extracted from the time-course analysis of the raw image data. Figure 4b displays the mean ‘on’ and ‘off’ times for each solution as a bar chart. Mean ‘off’ times were found to be significantly (ANOVA, p< 0.01) affected by the choice of electrolyte with values ranging between 0.1 s and 1 s. The variation in mean ‘on’ times across electrolytes was comparatively low (26 ms to 32 ms). Notably the mean ‘on’ and ‘off’ times differ by up to two orders of magnitude. This finding is in contrast to previous studies of NV photoblinking which report comparable on and off times however such studies consider single NVs and use higher temporal resolution than the 10 ms exposure time in the current work. Moreover, the detection of blinking events is contingent of sufficient image contrast between the on and off states which may further complicate detection of events, particularly over the 10 ms integration time used here.

**Fig. 4.**
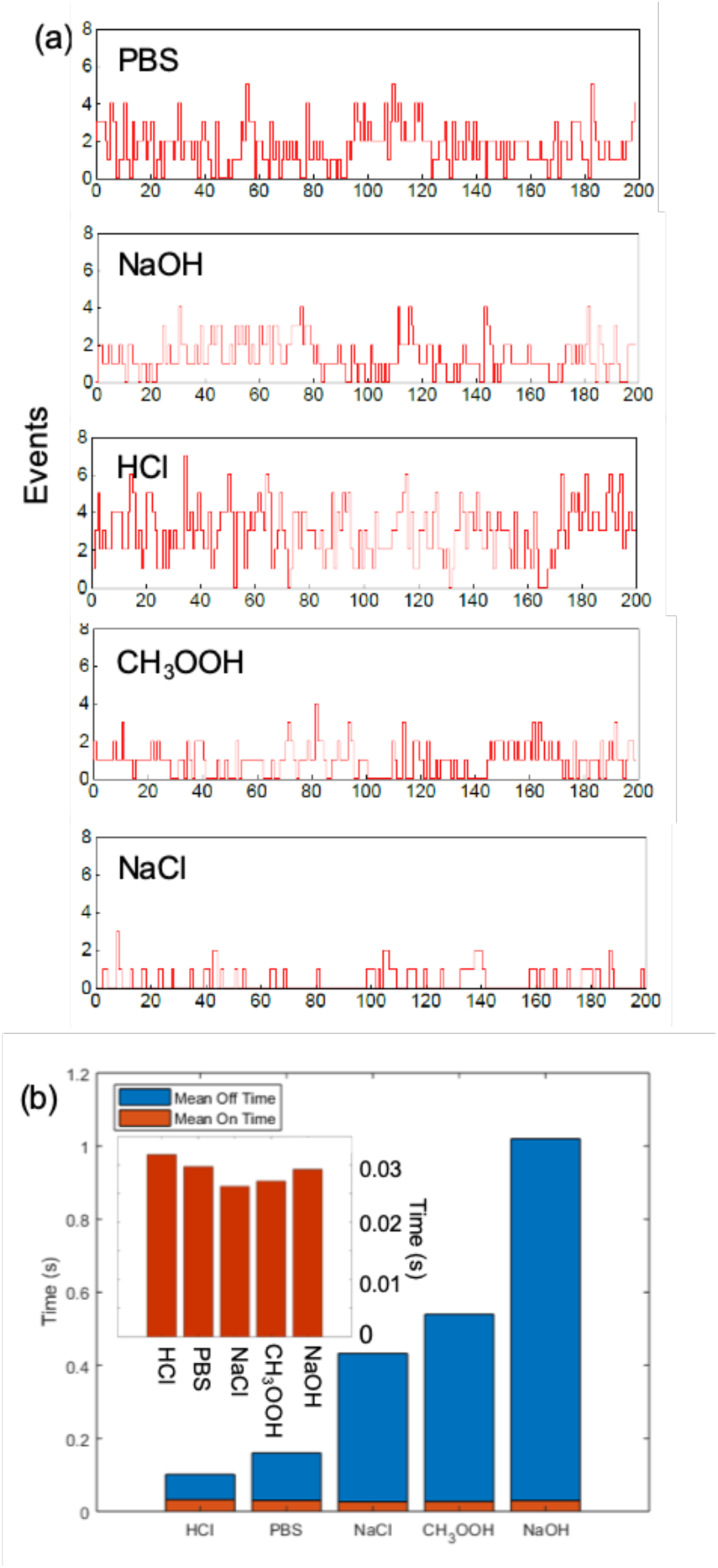
Effect of electrolyte medium on the temporal patterns of photoblinking in sandSTORM. (a)Time traces of blinking events per frame for 200 frames and (b) mean ON time (expanded detail) and OFF time for a single event in 5 different electrolyte solutions.

To investigate the temporal patterns of NV photoblinking and the underlying mechanisms, histograms depicting the probability distribution of ‘on’ and ‘off’ times were produced and fitted using a single exponential decay function, see Figure 5 and Supplementary Figure 2. The exponential nature of the probability distribution supports the hypothesis of charge state conversion owing to photoionization and is distinct from previous reports suggesting photoblinking is driven by electron tunneling and thermally dependent processes which characteristically follow a power law dependence. The exponent for each decay curve was also calculated. This revealed variations across the electrolytes used for both ‘on’ and ‘off’ times. Indeed, these differences were found to be at least an order of magnitude in the decay exponent between the corresponding ‘on’ and ‘off’ times for each electrolyte, readily apparent from the semi-log plots (Figure 5c and 5d). Collectively the experimental results suggest the chemical environment local to NVs strongly affects photoblinking and it is hypothesized that this may arise from a combination of effects including changes in radiative and non-radiative decay paths and lifetimes along with ligand induced shifts in the energies of the dark and bright exciton states [44]. It is further noted that blinking arising from charge state conversion between NV^-^ and NVovia two photon process, initiated by long term photoionization, is liable to be affected by the local chemical environment owing to inhibited recombination in the presence of charge traps near to and within the ND that prevents an efficient diffusion of charge carriers [43]. Moreover, electronic interactions between the NVs and surrounding molecules, including bond-building/breaking and charge transfer, will strongly affect the NV electronic states and correspondingly photoblinking [45].

**Fig. 5.**
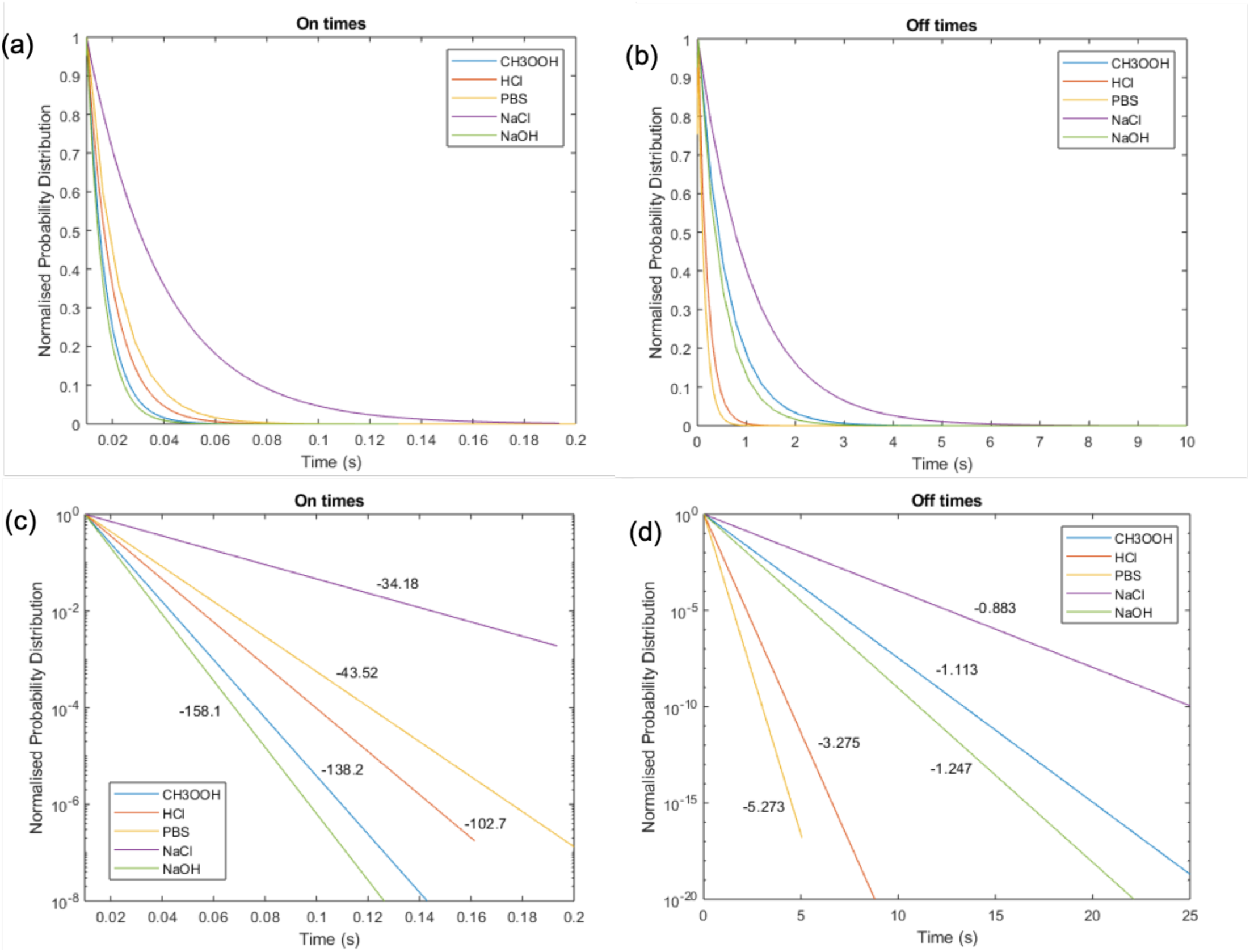
Temporal parameters of self-activated NV photoblinking in different electrolyte environments. Normalized probability distributions of (a) ‘on’-times and (b) ‘off’-times in CH_3_COOH, HCl, PBS, NaCl and NaOH, shown as fitted mono-exponential decay functions. (c & d) The respective probability distributions shown as semi-log plots.

Experiments presented in Figure 4a also revealed photoblinking could be dynamically adjusted by simply changing the solution in which NDs were imaged. Over the experimental time frame used here photoblinking was not observed from NDs under dry conditions, in freshly deionized water or in media containing significant proportions of glycerol. Further, photoblinking within PBS was found to be independent of oxygen scavengers or reducing agents, typically used in techniques similar to dSTORM. In summary, the data support an environmental-based sensitivity in photoblinking behaviour of NVs which could be further exploited to actively accelerate blinking and correspondingly reduce the duration of sandSTORM measurements.

### Comparison between sandSTORM and dSTORM

The usefulness of sandSTORM as a SMLM technique was evaluated through direct comparison with images of RyRs obtained using dSTORM, with particular emphasis on localization accuracy and image acquisition time. Figure 6 displays rendered dSTORM (Figure 6a) and sandSTORM (Figure 6b) images of RyRs from the same muscle cryosection. Qualitatively, the features seen in both images agree with the expected structural organization of RyR arrays as reported in previous dSTORM studies [37]. To quantitatively compare sandSTORM and dSTORM modalities, histograms were produced to assess the lateral event localization error (Figure 6c) along with Fourier Ring Correlation analysis as a measure of image resolution and as a means to assess the number of image frames and hence imaging time required to attain sub-diffraction resolution (Figure 6d). Figure 6c shows comparable lateral localization error for sandSTORM and dSTORM methods (modes of ∼ 18 nm and ∼ 20 nm respectively) but the range of localization errors for sandSTORM is markedly smaller than in dSTORM (95% of the events within a range of 32 nm and 62 nm) in similar samples. As a SMLM modality, the faster photoblinking of sandSTORM translates to quicker completion of the image of similar samples. Fourier Ring Correlation (FRC) values which, based on the implementation by Nieuwenhuizen, et al. [46], reports the momentary resolution of the image was used to characterize the temporal course of the image formation. In Figure 6d, the FRC value reached 95% of its minimum at 154 s with sandSTORM at an ∼ 3-fold speed advantage over the 465 s for dSTORM. The time course of reconstructing the geometrical features of the image was examined by plotting the Pearson correlation coefficient between the momentary image and the reconstruction from the final (30,000 frames) image (inset). 90% of the maximum correlation value was within 1000 s (16 min) for sandSTORM and ∼ 3000 s (50 min) for dSTORM. Collectively the results shown in Figures 6c and 6d demonstrate sandSTORM techniques enable attainment of super-resolution images significantly faster than dSTORM opening up the possibility for live cell imaging.

**Fig. 6.**
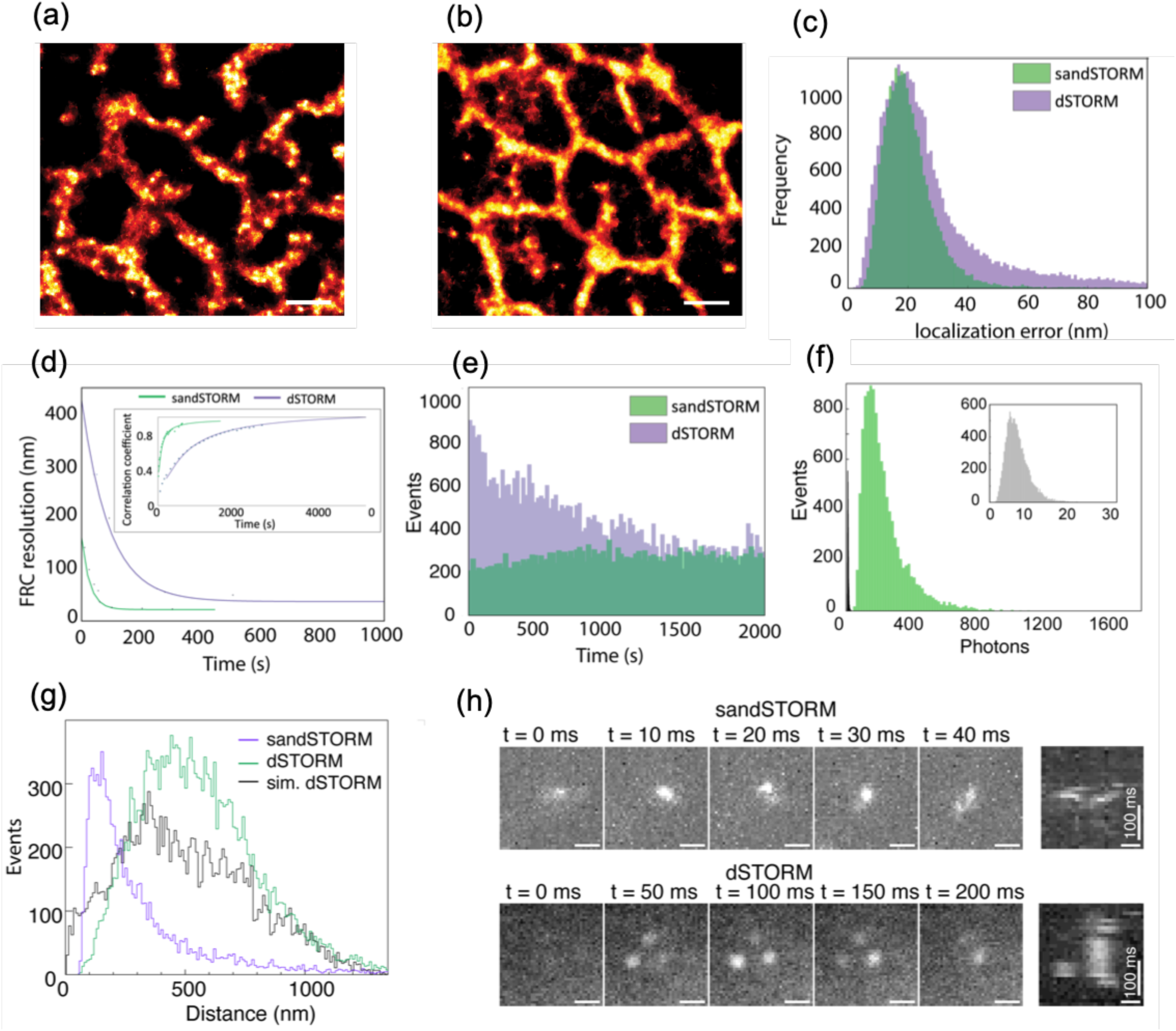
Comparison between sandSTORM and dSTORM. Rendered (a) dSTORM and (b) sandSTORM images of RyR labelling in fixed skeletal muscle tissue. Overlay plots show (c) histograms of the localization error of events, (d) temporal change in the resolution estimated with Fourier Ring Correlation (inset : time function of Pearson correlation coefficient relative to final image) and (e) temporal histogram of localization event counts for sandSTORM (green) and dSTORM (purple). (f) Overlaid frequency histograms of background photon count of dSTORM (green) and sandSTORM (grey; axes rescaled in inset). (g) Overlaid histograms of nearest neighbor distance for each event localized in a given frame in sandSTORM, dSTORM and events simulated to be spatially-random within the image frame (sim. dSTORM). (h) Serial frames of sandSTORM (upper row) and dSTORM (lower) image sequences illustrating the spatially-linked ND photoblink events. Panel on right shows corresponding x-t kymograph of the events through time (in vertical dimension). Scale bars: (a&b) 1 µm, (h) 400 nm

Photobleaching is a known common limitation of dSTORM that compromises localization accuracy over time owing to a reduction in the number of blinking events. Studies carried out herein demonstrate this limitation can be overcome through the use of NVs as fluorescent probes via the sandSTORM method. Experimental results are depicted in Figure 6e which enables a comparison between the number of blinking events recorded over time for sandSTORM and dSTORM. It is apparent that over the time period studied dSTORM suffered from photobleaching, a feature which depends heavily on the redox environment in the imaging field, whilst sandSTORM was resistant to such bleaching. This demonstrates a further strength of sandSTORM and the comparative suitability of sandSTORM for long-term or repeated imaging of samples as compared to dSTORM. As a further assessment of the two STORM methods the number of photons associated with individual blinking events was compared with the number of background photons in raw image frames.

The lower excitation power required for sandSTORM (less than two orders of magnitude compared to dSTORM; see Table 1) offers the added benefit of a background photon count which is ∼ 100-times lower in sandSTORM compared to that of dSTORM (Figure 6f). This avoids evoking of intrinsic autofluorescence which is a hallmark of large cells and optically-thick tissues and, proportionally reduces the contribution of out-of-focus photoblinking to preserve the contrast of the in-focus blink events. is the correspondence between the location at which the peak in the distribution for sandSTORM events occurs with ND size in Figure 6g. Differences in the spatial distribution of blinking events between the two methodologies is also seen from inspection of consecutive image frames for sandSTORM (Figure 6h; top row) and dSTORM (bottom row) and in the corresponding kymographs, depicting the fluorescent emission at

**Table 1.**
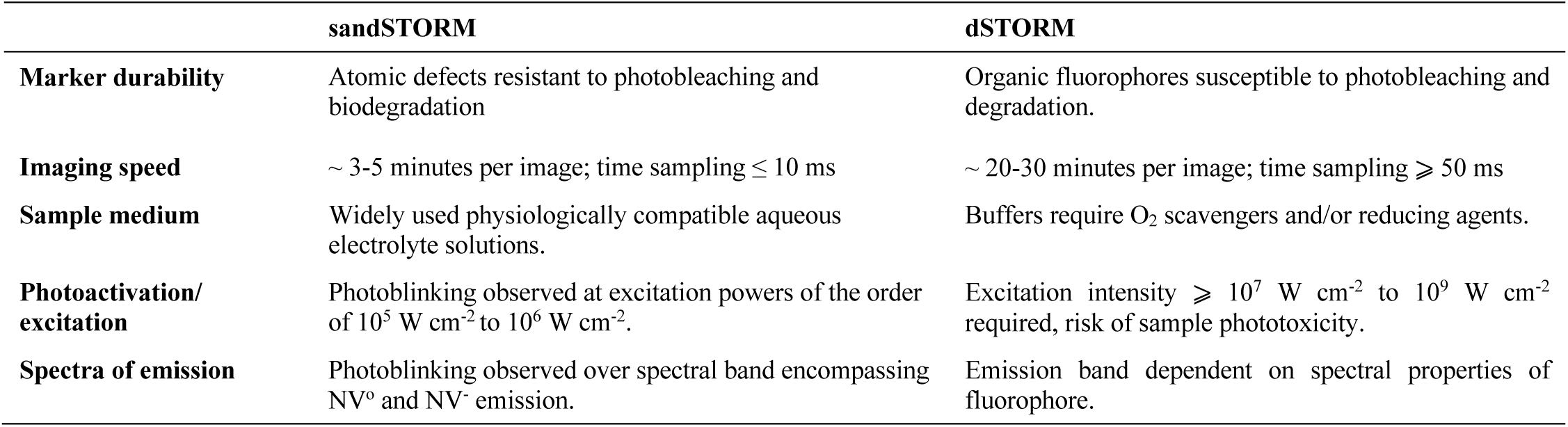
Comparison of key imaging characteristics of sandSTORM and dSTORM techniques.

|  | sandSTORM | dSTORM |
| --- | --- | --- |
| **Marker durability** | Atomic defects resistant to photobleaching and biodegradation | Organic fluorophores susceptible to photobleaching and degradation. |
| **Imaging speed** | ~ 3-5 minutes per image; time sampling $\leq 10$ ms | ~ 20-30 minutes per image; time sampling $\geq 50$ ms |
| **Sample medium** | Widely used physiologically compatible aqueous electrolyte solutions. | Buffers require $O_2$ scavengers and/or reducing agents. |
| **Photoactivation/excitation** | Photoblinking observed at excitation powers of the order of $10^5$ W $cm^{-2}$ to $10^6$ W $cm^{-2}$. | Excitation intensity $\geq 10^7$ W $cm^{-2}$ to $10^9$ W $cm^{-2}$ required, risk of sample phototoxicity. |
| **Spectra of emission** | Photoblinking observed over spectral band encompassing $NV^0$ and $NV^-$ emission. | Emission band dependent on spectral properties of fluorophore. |

The spatial distribution of blinking events was also investigated for both sandSTORM and dSTORM. In typical dSTORM experiments, the spatial distribution of blinking events associated with the aromatic fluorophores used within a given field of illumination, assuming uniformity of illumination, is random. It is hypothesized here that the physical size of individual NDs within a field of illumination will have a bearing on the spatial distribution of blinking events. To investigate this a histogram analysis of the nearest-neighbor distances between events localized within the same image frame was performed for sandSTORM and dSTORM data (Figure 6g) using a multi-emitter localization protocol optimized specifically for STORM data acquired with sCMOS cameras [47]. The histograms demonstrate a broader distribution of nearest neighbors for dSTORM as compared to sandSTORM. Of particular note fixed positions in the image frame over time.

Finally, a comparison of the characteristics of sandSTORM and dSTORM techniques is given in Table 1. This comparison highlights the advantages of sandSTORM particularly in relation to imaging speed, compatibility with existing biological protocols and suitability for long term imaging within live cells.

## Conclusion

Herein, a method for rapid SMLM that harnesses the self-activated photoblinking of NVs within NDs has been demonstrated. Specifically, the newly introduced method, sandSTORM, has been successfully applied to the study of two distinct and important biological systems, namely membrane receptors within tissue sections and networks of neuronal cells. The observed photoblinking of NV emission was concluded to be driven primarily by photoionization as a result of continuous illumination of samples with light from a 561 nm laser. Owing to the observed kinetics and spectral dependence of blinking it is considered that conversion from the NV° to the NV^-^ charges state is the dominant effect being observed alongside a two-photon initiated conversion from the NV^-^ to the NV° charge state. Interestingly it was shown that blinking could be accelerated through changes in the electrolyte in the buffer solution the biological sample was imaged in. This manifests itself as a reduction in blinking off time and is thought to arise as a result of electronic interactions between the NVs and surrounding molecules, including bond-building/breaking and charge transfer.

Major advantages of sandSTORM over existing SMLM methods include the high frame rates super-resolution localization can be achieved at (100 fps (sandSTORM) compared with 0.1 fps (dSTORM)), the simple protocols for incorporation of NDs within samples, the readily accessible experimental implementation within commercial fluorescent microscope and the temporal stability of photoluminescence enabling long term and replicate studies to be performed. Overall, sandSTORM is a SMLM technique with the potential to transform super-resolution imaging within biological systems with particular applications in live cell imaging owing to the fast imaging times, biologically favourable properties of NDs and compatibility of the experimental protocols with existing microscopy platforms in wide use within biological settings.

## Materials and Methods

### Immunohistochemistry

The extensor digitorum longus (EDL) muscles were dissected from fresh carcasses of healthy male Wistar Crl rats weighing ∼ 200 g euthanized for heart tissue dissection in Prof Derek Steele’s laboratory. Muscles were dissected with intact tendons before pinning out in standard Ringer’s saline solution. The muscles were immediately fixed in 2% paraformaldehyde (PFA; w/v) in phosphate buffered saline (PBS) for 1 hour at 4 °C. Fixed muscles were washed in fresh PBS for 10 minutes and cryoprotected in PBS containing 30% sucrose. They were frozen in methyl butane cooled in liquid nitrogen, mounted onto a Leica CM 1900 cryostat set to −25 °C using OCT compound (Tissue-Tek, USA) and cut into 10 µm thick cryosections. Sections were mounted onto number 1.5 glass coverslips (Menzel-Glaser; Germany) coated with 0.05% (w/v) poly-L-lysine (Sigma-Aldrich, MO).

Coverslips were attached to the underside of custom-made acrylic stage adapters using silicone elastomer Pinkysil (Barnes, New Zealand) such that an open chamber is formed with the tissue section was positioned in it. Prior to staining, the sections were briefly hydrated and then blocked with Image-iT FX Signal Enhancer (Thermo Fisher, MA) for 1 hour at room temperature (∼ 21°C). The incubating the primary antibodies in the antibody incubation buffer overnight at 4°C. Coverslips and sections were rinsed in fresh PBS in three 10-min steps prior to the application of the secondary antibodies for 2 hours at room temperature. Three further rinsing steps were applied prior to immersing the sections in the imaging buffer.

### ND conjugation to secondary antibodies

100 ul of NDs of 50 or 100 nm (1 mg/mL) were dispersed in 400 ul 1M HEPES buffer at pH 5. Ethyl dimethylaminopropyl carbodiimide (EDC; Sigma; 0.04 M) and N-hydrosuccinimide (NHS; Sigma; 0.01 M) was added into the ND suspension and sonicated for 1 hour in a sonicator bath. The suspension was centrifuged at 18 000 rcf for 10 minutes to pellet NDs and the supernatant was discarded. Centrifugation was repeated with 1x PBS three times to wash NDs. 500 ul of PBS was added and sonicated to disperse ND pellet. 20 ul of secondary antibody (AffiniPure goat anti-mouse IgG or AffiniPure goat anti-rabbit; Jackson Immuno Research, USA) was added and incubated at 4oC overnight. The suspension was then centrifuged at 18 000 rcf for 10 minutes at 4oC and washed with PBS once. The suspension was dispersed in 400 ul of PBS and 0.05% sodium azide (Sigma).

All animal studies were conducted in compliance with the ethics and animal welfare in place in the University of Nottingham, in accordance to the 1986 Animals (Scientific Procedures) act. 128

### Neuronal cell culture

C57/BL6 mouse embryos in E16-E17 developmental stage were culled and their brains removed. The brain cortices were dissected, and the meninges separated under a dissection microscope. The tissue was further incubated in Hanks Balanced Salt Solution (HBSS, Ca2+ and Mg2+-free; Gibco) with 1mg/ml trypsin and 5mg/ml DNAseI (both Sigma) at 37oC/5% CO2 for 30 minutes. Following treatment with 0.05% (v/v) trypsin inhibitor (Life Technologies), the tissue was washed in Neurobasalmedia (Gibco) and 5mg/ml DNAseI was added before mechanical dissociation of the tissue. The cells were finally washed in Neurobasal media and spun down at 250x g for 5 minutes. Dissociated cells were further resuspended in Neurobasal media supplemented with 1x GlutaMax and 2% (v/v) B-27 (both Gibco) and seeded in 35mm μ-Dishes high (Ibidi) with or without diamond plate at 1.25×105/cm2. Neuron cortical networks were allowed to develop for 14 days before incubation with the NDs. 100nm red fluorescent NDs (fND biotech) were added to cultures at a concentration of 30ug/ml to allow cell-mediated endocytosis of the NDs.

Cortical neurons containing NDs were fixed 24 hours later in 4% (w/v) paraformaldehyde (3.6% sucrose (w/v), 1x PBS, 5mM MgCl2, pH 7.4; ThermoFisher) for 30 minutes at room temperature, washed with 10mM Glycine in PBS and further permeabilized in PBS/Glycine-Triton (1x PBS, 10mM glycine, 0.2% (v/v) Triton X-100; Sigma) for 20 minutes. The cultures were then blocked for 1 hour at room temperature with 3% (w/v) BSA in PBS (Sigma), followed by incubation with anti-Gfap (1:100; Abcam) and anti-acetylated tubulin (1:300; clone 611B-1, Sigma-Aldrich) in blocking buffer at 4oC/overnight. After PBS-Triton 0.1% (v/v) washes, secondary antibodies Alexa Fluor 488 (1:300; Molecular Probes) were incubated where appropriate for 1 hour.

### Materials and solutions

Red fluorescent NDs of ∼ 50 nm diameters with a population of NV0 and NV-were used for these experiments (Catalogue number brFND-100; FND Biotech, Taiwan). The Ringer solution used for dissecting the fresh muscle tissue was designed previously [48], and contained 112 mM of NaCl, 3.3 mM of KCl, 2.5 mM of CaCl2, 1 mM of MgCl2 and 20 mM of HEPES with the pH adjusted to 7.4 using NaOH. Primary and secondary antibodies were incubated in PBS containing 0.05% NaN3, 2% bovine serum albumin (Sigma), 2% normal goat serum (Sigma) and 0.05% Triton X100 (Sigma). The imaging buffer for dSTORM contained 90% Glycerol (v/v; Sigma) and 15 mM β-mercaptoethylamine (Sigma) in 1xPBS at pH 8.1.

A range of aqueous imaging solutions were teste d for sandstorm. Four ionic solutions were prepared in distilled deionized water which were HCl and CH3COOH solutions were prepared at pH 3.45, NaOH at pH 10.55, and NaCl at a concentration of 0.35 mM. DD water and PBS 1x (Sigma) were also used.

Primary antibodies used for immunohistochemistry were mouse monoclonal anti-RyR raised against partial RyR1 of chicken pectoral muscle [49]; MA3-925; Thermo Scientific), Alexa Fluor 680-conjugated highly cross-adsorbed (H+L) goat anti-mouse IgG and goat anti-rabbit IgG antibodies were used as secondary antibodies in dSTORM labelling experiments.

### Image acquisition

Samples were imaged on a modified Nikon TE2000 inverted microscope with a Nikon 1.49NA oil-immersion TIRF objective which focused excitation light onto the sample in HiLo configuration [50]. This allowed the illumination of ∼ 8 µm wide approximately circular area in-plane and ∼ 5 µm-deep volume. For imaging Alexa 680 markers (dSTORM), the laser beam from a solid-state 642 nm laser (Cobalt, Sweden) was focused onto the sample at a power of ∼ 2.0 × 10^8^ W cm^-2^. Emission light was passed through a Q690LP dichroic mirror (Chroma Technology) and an DC/ET720/60m emission filter (Chroma) and recorded onto a Zyla 5.5 USB3.0 scientific-CMOS camera (Andor Ltd, Belfast) at an integration time of 50 ms/frame. For sandSTORM, A 561 nm 200 mW solid-state laser (CNI lasers, China) was used in combination with an AT565DC dichroic mirror (Chroma). Emission filters used for sandSTORM were the 605/70 nm 49004ET-Cy3/TRITC (Chroma), 692/40 nm BrightLine (Semrock), HQ520/40 nm (Chroma), 575/5 nm (Semrock), and 590 nm LP (Brightline) Image acquisition for sandSTORM was set at 10 ms/frame integration. The power of the 561 nm excitation at the focal plane was measured to be between 5.0 × 10^5^ and 1.0 × 10^6^ W cm^-2^. To alter the effective excitation intensity, the laser beam was passed through a motorized set of neutral density filters (Thorlabs; Germany).

### Image analysis and reconstruction

Single molecule events in each image frame were detected using a multi-threshold detection algorithm. Each detected event was localized by fitting the event with an adaptive two-dimensional Gaussian model using a least-squares fitting procedure adopted in an algorithm described previously [51]. These algorithms were implemented in freely available Python Microscopy Environment (PyME) software (freely available via www.python-microscopy.org) concurrent to image acquisition on a Lenovo ThinkTower workstation with a quad-core CPU, 16 Gb of memory and a 1 Tb SSD. Localized coordinates, time stamps and single molecule fitting parameters were saved in HDF format which allowed statistical analysis of parameters such as the localization error and event localization rate.

Greyscale 32-bit TIFF images encoding local densities of localized events were rendered using a protocol based on Delaunay triangulation [52]. Here, each detected event coordinate was jittered randomly in 3D at an amplitude that was proportional to the average distance between this point and its nearest neighbor prior to the calculation of Delaunay triangulation. Averaging between ten independent triangulations produced a greyscale image at a pixel sampling of 5 nm/pixel with minimal background where intensity was proportional to the local event density.

The number of events per frame were analyzed using TrackMate in ImageJ [53]. Image stacks (40,000 frames, 10ms/frame, N=3) were processed in ImageJ, initially using a rolling ball background subtraction based on a 10-pixel kernel, followed by binary thresholding using the ‘minimum’ filter to allow visualizing of the ‘on’ blinking events against a zero baseline for the ‘off’ times. Nine to ten regions of interest were selected within each experimental repeat, corresponding to the location of NDs separated by a distance greater than the point spread function of the objective. A fluorescence time trace for region was plotted in OriginPro 9.0, and the traces were analyzed using the Peak Analyzer toolbox in Origin, to obtain the initial and end frame numbers for each detected peak (**SI Fig. 1)**. ON times were then calculated by tracking the full duration of each event where peaks were detected, whilst OFF times were calculated by subtracting the end frame value of each peak from the start frame value of the next peak in the trace. Mean on and off times were calculated by taking the average for each of the 9 regions across three separate experimental repeats (total 27-30 NDs).

Analysis of time evolution of resolution: For assessing the resolution at a given time point in the image acquisition, the localized coordinates of events were binned into alternating time blocks of 2000 frames up to the time point of interest. The alternating time blocks are combined into two independent sub-samples of event coordinates which were used for calculating the Fourier ring correlation, in PyME as described by [46]. The point at which the spatial frequency of the FRC intersects with 1/7 of the peak correlation was taken as the resolution at that time point.

Completion of the image acquisition, in addition to reaching the finest achievable resolution, requires optimal sampling of the marker population in the field of view. To assess this, the image rendered from the cumulative of the localized events up to each time point was subjected to a retrospective Pearson correlation with a final version of the rendered image, implemented in PyME. The time point at which the Pearson’s correlation coefficient exceeds 0.95 of the final image was taken as the ‘time to maximum correlation’.

### Nearest neighbor analysis and simulation of synthetic STORM data

dSTORM and sandSTORM events were localized within a 8 × 8 μm window using a custom-written multi-emitter localization algorithm described previously [54] and available in PyME. Two-dimensional coordinates and the time stamps (in image frame number) of the events localized with this routine were analyzed further in a custom-written script. Where each image frame recorded multiple localized events, the distance from the centroid of each event to that of its nearest neighboring concurrent event was recorded and plotted as frequency histograms.

To simulate a random blinking spatiotemporal pattern, a synthetic image series was rendered via PyME (as described previously in the supplementary section of [7]) using the rendered sandSTORM images as a starting map of spatial density of markers. The localization and image acquisition parameters (e.g. localization error, sigma of each fit, amplitude and series length) were set to mimic dSTORM experimental data. The simulation parameters included pre-set values such as average event number per pixel of the starting image (1.0), mean event intensity (1000 photons), mean event background intensity (200 photons), mean event duration (150 ms), mean number of detections per marker (2.0) and number of image frames (30,000).

### Statistical analysis

Statistical analysis was conducted using OriginPro 9.0. A one-way ANOVA statistical analysis was performed for the ON and OFF times for events (N=3). P values were denoted as *p<0.05 to indicate statistical significance between groups.

## End Matter

### Supporting information

The authors wish to dedicate this publication to Dr Simon Levett, our beloved colleague and friend, who sadly recently passed away. Simon was a fundamental member of our research team who was a source of constant support and scientific insight.

Supporting Information is available from the figure supplement or from the author.

### Author Contributions and Notes

Experiments were conceived by K.K.N., I.J., J.C.P. & M.L.M. Experiments were performed, and data were acquired by K.K.N., J.C.P & R.M-R.. Data were analyzed by K.K.N., I.J., J.C.P. & MLM, interpreted by K.K.N., I.J., J.C.P. & M.LM. The manuscript was written by K.K.N., I.J, M.L.M & J.C.P.

The authors declare no conflict of interest.

## Acknowledgments

The authors thank the Wellcome Trust (Seed Award awarded to IJ; Ref 207684/Z/17/Z awarded to IJ) and the European Research Council (ERC) (ERC Consolidator Award, TransPhorm, grant number 23432094 awarded to MLM) for the funding which supported this research. Also acknowledged, are Tim Munsey for his assistance with dissecting and sectioning the muscle tissue used for imaging experiments, Dr David Baddeley (University of Auckland) and Prof Christian Soeller (University of Exeter) for helpful discussions on the adaptation of PyME for the image analysis presented in this manuscript and Drs David Simpson and Liam Hall (University of Melbourne) for useful discussion on NV charge state dynamics.

