## Supplementary figures for "Self-activated photoblinking of nitrogen vacancy centers in nanodiamonds (sandSTORM): A method for rapid single molecule localization microscopy with unlimited observation time"

### Slide 1
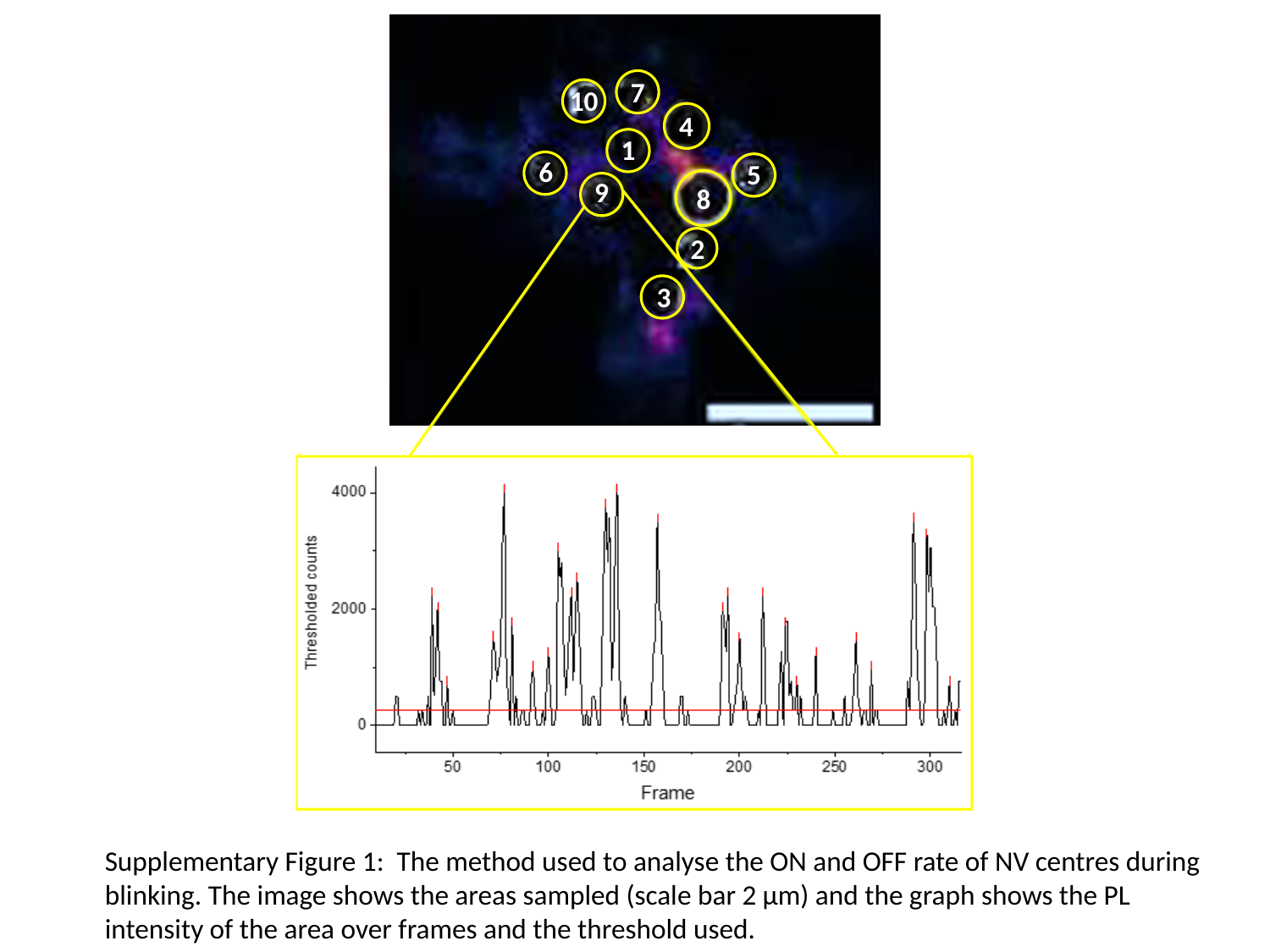

7
10
4
1
6
5
9
8
2
3
Supplementary Figure 1: The method used to analyse the ON and OFF rate of NV centres during blinking. The image shows the areas sampled (scale bar 2 µm) and the graph shows the PL intensity of the area over frames and the threshold used.

### Slide 2
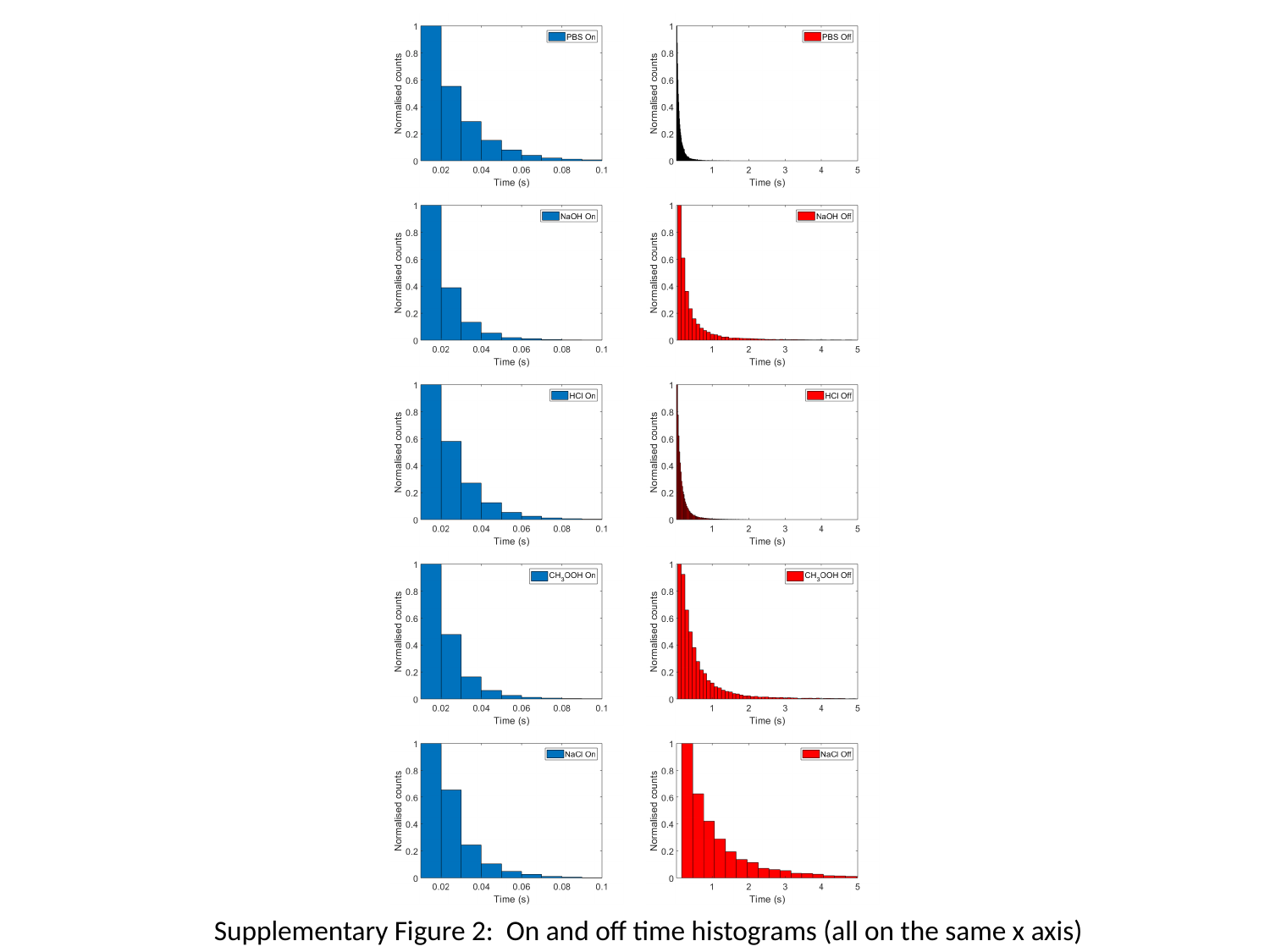

Supplementary Figure 2: On and off time histograms (all on the same x axis)

### Slide 3
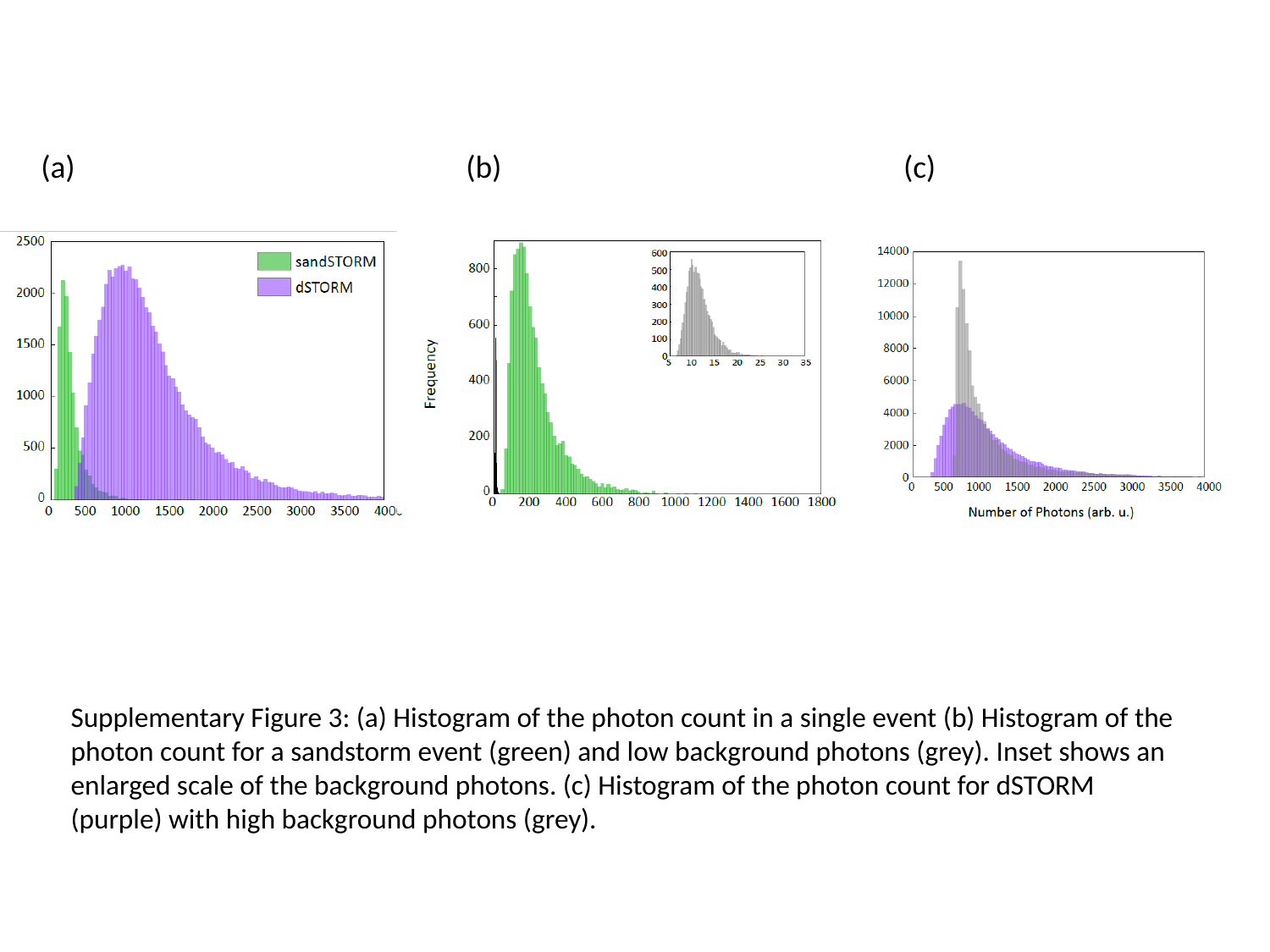

(a)
(b)
(c)
Supplementary Figure 3: (a) Histogram of the photon count in a single event (b) Histogram of the photon count for a sandstorm event (green) and low background photons (grey). Inset shows an enlarged scale of the background photons. (c) Histogram of the photon count for dSTORM (purple) with high background photons (grey).
