## Supplementary movie for "Self-activated photoblinking of nitrogen vacancy centers in nanodiamonds (sandSTORM): A method for rapid single molecule localization microscopy with unlimited observation time"

### Slide 1
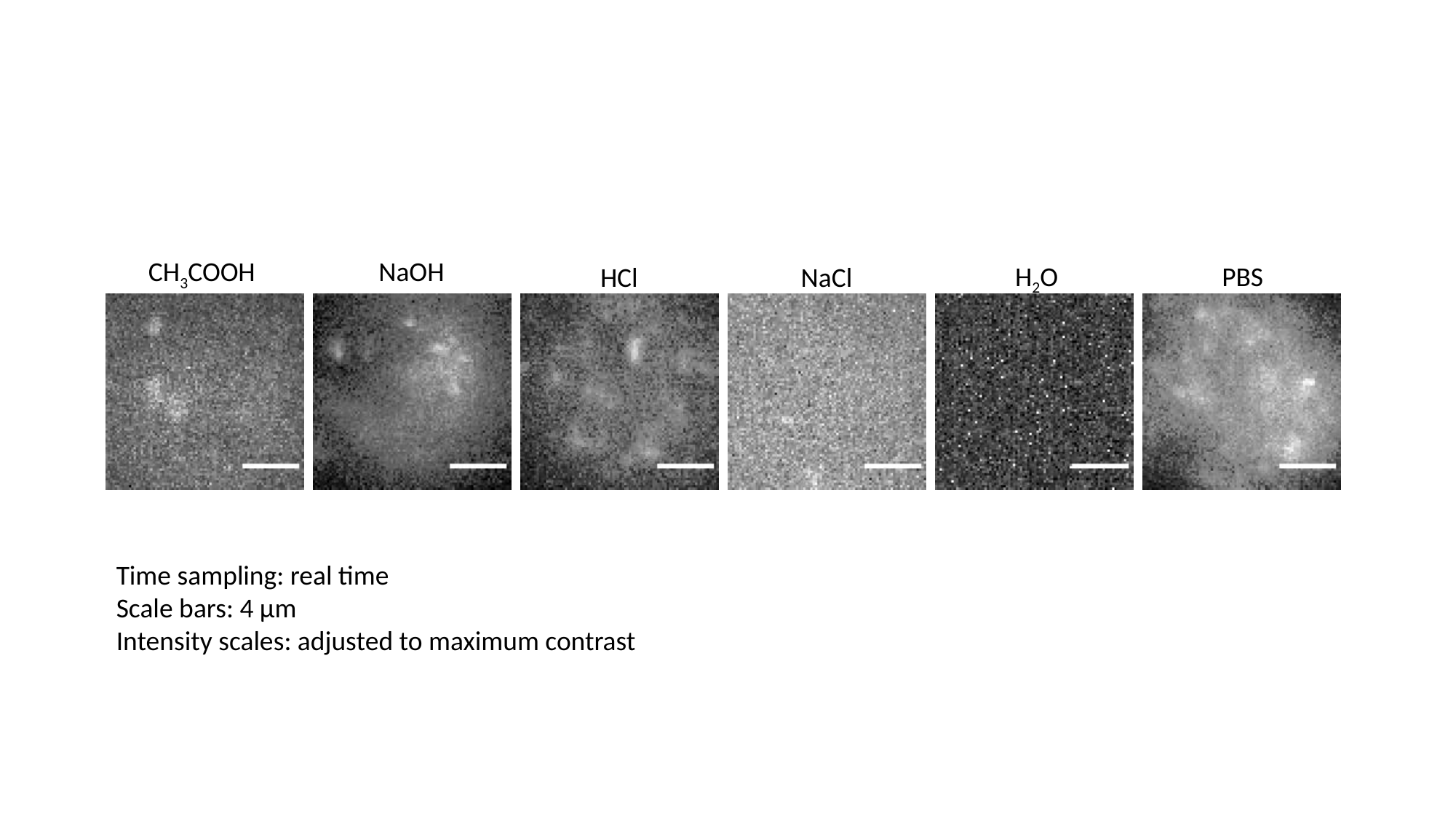

CH_3_COOH
NaOH
H_2_O
PBS
HCl
NaCl
Time sampling: real time
Scale bars: 4 µm
Intensity scales: adjusted to maximum contrast
